# Hybrid Mechanistic–Neural Modeling of Concentration-Time Dynamics Generalizes Human Pharmacokinetics Prediction Across Unseen Chemical Space

**DOI:** 10.64898/2026.10.01.756018

**Authors:** Amitesh Badkul, Lianjin Cai, Minseok Kim, Junmei Wang, Lei Xie

**Author notes:** Contributing authors. These authors contributed equally to this work (Lianjin Cai, Minseok Kim).

## Abstract

The pharmacokinetics concentration-time profile encodes ADME dynamics. Its optimization in drug discovery curbs late-stage attrition, yet forecasting human pharmacokinetics from chemical structure remains difficult. Machine learning models ignore continuous ADME dynamics, whereas physiologically based pharmacokinetics (PBPK) models demand unavailable parameters and impose rigid, misspecified compartmental structures. We present PK-MUSE (**P**harmaco**K**inetic hybrid **M**echanistic **U**niversal **S**ystem **E**quation), augmenting a two-compartment model with bounded, state- and time-dependent neural corrections to elimination and intercompartmental distribution. Molecular structure parameterizes compound-specific behavior, while learned corrections refine dynamic equations. Across scaffold and temporal out-of-distribution evaluations, PK-MUSE achieved the lowest mean exposure-endpoint MdFE, reducing MdFE by 7.2%, 45.2%, and 26.2% versus state-of-the-art machine learning (HML-RF), deep learning compartmental (DeepCt), and PBPK (PK-Sim) models. Interrogating the corrections localized misspecification to distribution rather than elimination. By extending mechanistic simulators with data-driven constrained neural dynamics that preserve interpretability and generalize to unseen chemotypes, PK-MUSE advances digital twins for drug discovery.

## 1 Introduction

Drug discovery and development frequently fails, and poor pharmacokinetics (PK) remains a leading cause of clinical failure [1–3]. A PK profile describes how drug concentration changes over time after dosing, governed by absorption, distribution, metabolism, and excretion (ADME). It is a critical determinant of drug efficacy and safety, informing candidate selection, dose design, safety-margin assessment, and patient-specific PK modeling [1, 4, 5]. Reliable prediction of human exposure for new chemotypes is therefore valuable early in discovery, before experimental human PK data are available, because poor disposition can drive costly downstream failure [2, 6]. Yet human PK profile prediction remains a difficult structure-to-dynamics problem. Unlike most molecular machine learning (ML) tasks, which map a chemical structure to a scalar label such as solubility, clearance, or toxicity, PK profile prediction must output a time-indexed concentration curve whose shape depends on dose input, distribution, and elimination. This places it closer to dynamical-system learning than to standard property prediction. Human PK datasets also contain far fewer profiles than typical molecular property tasks, and this limited supervision makes it harder for flexible ML models to learn profile dynamics and generalize to new compounds.

Existing approaches address different sides of this problem. Classical compartmental and physiologically based pharmacokinetics (PBPK) models represent drug disposition with ordinary differential equations (ODEs), providing an explicit dynamical system and allowing concentrations to be propagated under different dosing conditions. Compartmental models reduce the body to a few mathematical compartments governed by rate parameters without direct physiological meaning, whereas PBPK models treat anatomical organs as compartments connected by blood flow, requiring a much larger set of compound- and system-specific parameters such as solubility, permeability, and intrinsic clearance [7]. Both remain useful because they encode pharmacological structure and support extrapolation [8, 9]. However, compartmental models can oversimplify drug behavior, while PBPK models demand many measured ADME parameters [5, 6, 10], giving rise to two central problems: model misspecification and parameter uncertainty. Misspecification arises when the assumed number of compartments or the fixed rate structure does not match a compound’s true disposition kinetics. A two-compartment model, for example, can misspecify the distribution phase when a compound exhibits multi-exponential behavior requiring three or more compartments, and this bias propagates into the predicted plasma concentration-time profile and associated systemic exposure metrics [11].

ML can assist PK profile prediction both explicitly, by predicting the profile directly, and implicitly, by predicting the parameters required as input to PBPK models, reducing dependence on measured values by learning PK endpoints from molecular structure [5, 12]. However, such parameter-prediction models can propagate errors into downstream simulations [13–15]. Pointwise concentration models can fit observed time points but do not learn a single dynamical process linking the profile across time [16, 17], and therefore lack the temporal consistency and pharmacological constraints that mechanistic models supply. This limitation is particularly relevant in model-informed drug development, where regulatory frameworks emphasize model credibility with adequate characterization of PK profiles. Sequence models such as LSTMs can represent ordered concentration measurements, but are typically tied to the observed sampling grid and struggle with sparse, irregular time points [18].

Scientific machine learning (SciML) is a relevant approach for data-driven learning in dynamical systems. It combines mechanistic models with data-driven neural networks, embedding trainable functions into otherwise structured dynamical models [19, 20] and thereby retaining physiological structure and a constrained solution space while still learning from data. Within this framework, neural ordinary differential equations (NODEs) replace a discrete sequence of neural network transformations with a learned continuous-time vector field integrated numerically by an ODE solver [21]. A complementary formulation, universal systems equations (USE, also known as universal differential equations), retains known mechanistic equations and introduces trainable functions only where the prescribed dynamics are insufficient [20]. NODEs and USEs have been applied to several PK problems, but mainly in settings distinct from de novo molecule-to-profile prediction [22]. Lu et al. modeled patient-level PK for a single therapeutic, using first-cycle measurements to predict subsequent cycles and extrapolate between regimens [22]. Valderrama et al. retained a one-compartment model and trained a network to learn an incompletely specified absorption process from simulated data while estimating distribution and elimination parameters [19]. Bräm et al. examined NODEs for fitting predefined PK systems and simulating changes in dose and administration [23]. These studies show that neural components can learn missing or complex PK dynamics, but none addresses the problem considered here: predicting the continuous human PK dynamics of an unseen compound from molecular information alone, without prior PK/ADME measurements for that compound.

Here we introduce PK-MUSE, a PK mechanistic-neural universal systems equation model that predicts human intravenous concentration–time profiles from chemical structure alone. PK-MUSE first maps molecular information to compound-specific PK quantities that parameterize a two-compartment mechanistic ODE, then augments these dynamics with bounded, state- and time-dependent neural corrections to three pharmacologically identifiable rate processes: elimination, central-to-peripheral distribution, and peripheral-to-central redistribution. The resulting effective rates remain positive and act within the original compartmental flux structure, adding learned flexibility without replacing the mechanistic backbone with an unrestricted neural vector field. We evaluate PK-MUSE under random, scaffold, and temporal splits that probe progressively harder generalization beyond the training chemical space, and compare it with state-of-the-art approaches spanning pointwise ML, deep learning compartmental prediction, and PBPK simulation. PK-MUSE consistently improves prediction across these regimes, with the learned corrections most beneficial when extrapolating to chemically distinct compounds. Crucially, the corrections remain interpretable in terms of named PK processes: they identify intercompartmental distribution, particularly during terminal disposition, as the dominant source of misspecification in the two-compartment model. PK-MUSE thus provides both a predictive framework for molecular PK and a data-driven means of locating where established mechanistic assumptions require refinement.

## 2 Results

### 2.1 Overview of PK-MUSE

PK-MUSE predicts human intravenous concentration-time profiles from molecular structure by combining a two-compartment PK ODE with bounded, state-dependent neural corrections to its three rate processes (Fig. 1A). The mechanistic backbone enforces mass-balance and positivity constraints that guarantee physically plausible trajectories, while the learned corrections add flexibility to elimination and inter-compartmental distribution rates without replacing the underlying compartmental structure. We evaluated PK-MUSE against its uncorrected mechanistic backbone (PK-MUSE_Mech_), a pointwise ML model (HML-RF), a deep learning compartmental model (DeepCt), and a PBPK simulator (PK-Sim) under three split strategies: a predefined random split, a scaffold split excluding entire Bemis-Murcko scaffolds from training, and a temporal split separating compounds by year of disclosure (pre- and post-1997). Performance was assessed with exposure endpoints, area under the curve (AUC) and the maximum concentration (*C*_max_) of a concentration-time curve, supplemented by a physiological-validity assessment that flags predictions with negative concentrations, rising terminal slopes, or sustained post-infusion rises.

**Fig 1.**
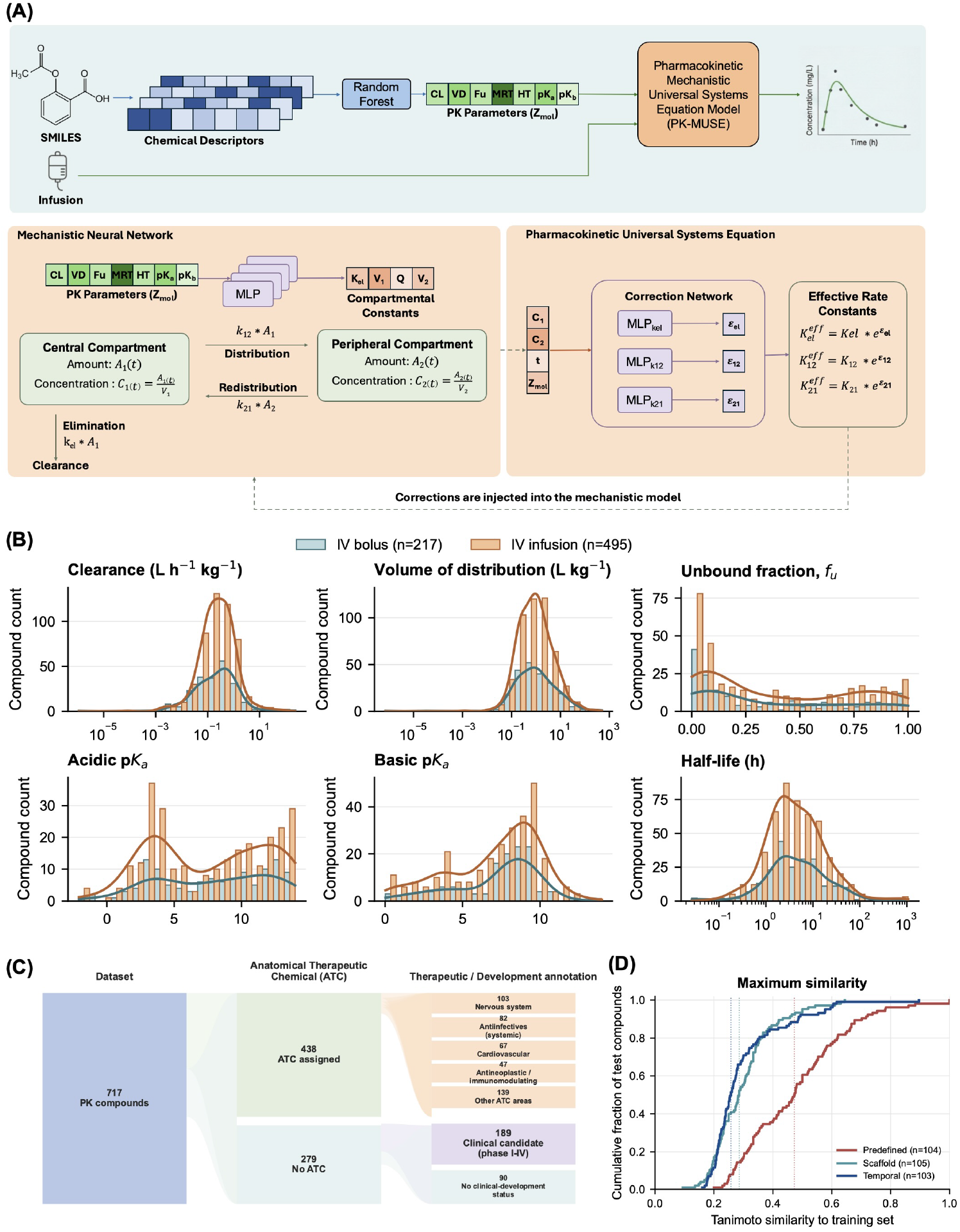
(A) Overview of the PK-MUSE framework. (B) Distributions of PK parameters present in the dataset by intravenous bolus and infusion administration. Bars represent compound counts, with overlaid kernel density estimates. (C) Dataset composition by Anatomical Therapeutic Chemical (ATC) assignment. (D) Empirical cumulative distributions of each test compound’s maximum Tanimoto similarity to the corresponding training set for the predefined, scaffold and temporal splits. Lower similarities indicate greater structural separation between training and test compounds.

Figures 1B–D characterize the dataset used in the study. Each of the PK parameters spans several orders of magnitude or the full pKa range, reflecting substantial pharmacokinetic diversity within the dataset. Panel C summarizes the dataset’s regulatory and therapeutic annotation. Out of the 718 compounds, 438 carry an assigned Anatomical Therapeutic Chemical (ATC) code [24] (predominantly nervous-system, antiinfective, cardiovascular, and antineoplastic/immunomodulating agents) and 279 do not, of which 189 nonetheless reached at least one clinical trial phase and 90 have no recorded current clinical development status. The PK properties of both these groups are not statistically different. Panel D shows, for each split, the empirical cumulative distribution of each test compound’s maximum Tanimoto similarity to its corresponding training set, computed on Morgan fingerprints (radius 2, 2048 bits). The scaffold and temporal test sets are shifted toward lower similarity than the predefined split, confirming that they impose a harder generalization test to less structurally similar compounds.

### 2.2 PK-MUSE generalizes across out-of-distribution chemical space

PK-MUSE consistently predicted clinically relevant exposure endpoints across the predefined, scaffold, and temporal regimes better than the baselines (Table 1). On the predefined split, it achieved an AUC median fold error (MdFE) of 1.925 and a *C*_max_ MdFE of 2.031, compared with 3.244 and 3.013 for DeepCt and 3.001 and 2.223 for PK-Sim. HML-RF was competitive in AUC prediction, with an AUC MdFE of 1.946 and an AUC *R*^2^ of 0.847 compared with 0.845 for PK-MUSE. However, PK-MUSE achieved lower AUC MdFE, higher twofold and threefold accuracy, and fewer invalid profiles than HML-RF.

**Table 1.**
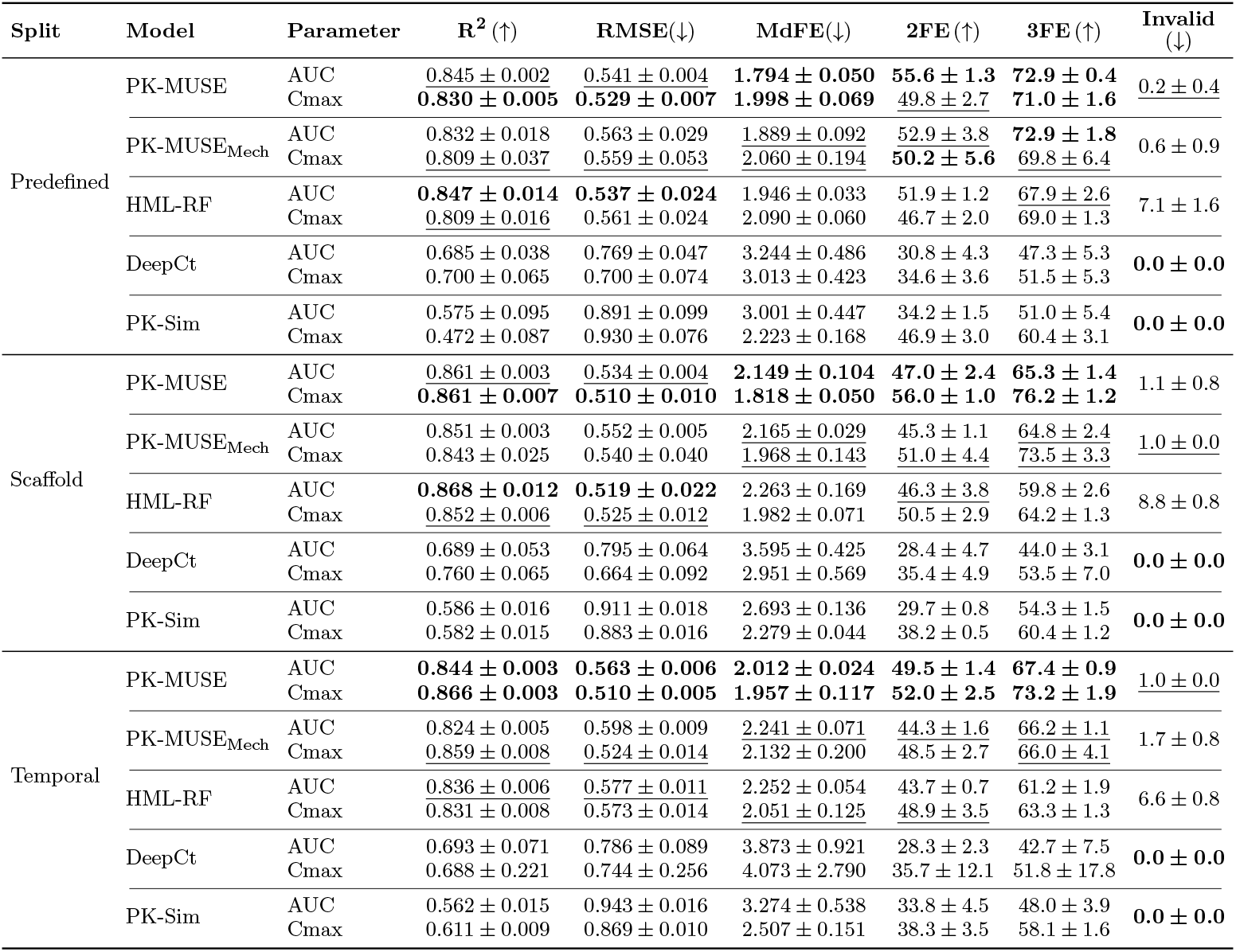
Exposure metric performance across split strategies. Best values are bold and second-best distinct values are underlined within each split and parameter. Invalid profiles are reported once per model because validity is a property of the predicted curve and does not depend on the exposure endpoint. *R*^2^ and RMSE were calculated from finite numerical endpoint predictions, whereas MdFE, twofold accuracy, and threefold accuracy assigned physiologically invalid profiles worst-case fold-error values.

| Split | Model | Parameter | $R^2$ (↑) | RMSE (↓) | MdFE (↓) | 2FE (↑) | 3FE (↑) | Invalid (↓) |
| --- | --- | --- | --- | --- | --- | --- | --- | --- |
| Predefined | PK-MUSE | AUC | <u>0.845 ± 0.002</u> | <u>0.541 ± 0.004</u> | **1.794 ± 0.050** | **55.6 ± 1.3** | **72.9 ± 0.4** | <u>0.2 ± 0.4</u> |
|  |  | Cmax | **0.830 ± 0.005** | **0.529 ± 0.007** | **1.998 ± 0.069** | **49.8 ± 2.7** | **71.0 ± 1.6** |  |
| | PK-MUSE<sub>Mech</sub> | AUC | 0.832 ± 0.018 | 0.563 ± 0.029 | <u>1.889 ± 0.092</u> | <u>52.9 ± 3.8</u> | **72.9 ± 1.8** | 0.6 ± 0.9 |
|  |  | Cmax | <u>0.809 ± 0.037</u> | <u>0.559 ± 0.053</u> | <u>2.060 ± 0.194</u> | **50.2 ± 5.6** | <u>69.8 ± 6.4</u> |  |
| | HML-RF | AUC | **0.847 ± 0.014** | **0.537 ± 0.024** | 1.946 ± 0.033 | 51.9 ± 1.2 | <u>67.9 ± 2.6</u> | 7.1 ± 1.6 |
| | | Cmax | <u>0.809 ± 0.016</u> | 0.561 ± 0.024 | 2.090 ± 0.060 | 46.7 ± 2.0 | <u>69.0 ± 1.3</u> | |
| | DeepCt | AUC | 0.685 ± 0.038 | 0.769 ± 0.047 | 3.244 ± 0.486 | 30.8 ± 4.3 | 47.3 ± 5.3 | **0.0 ± 0.0** |
| | | Cmax | 0.700 ± 0.065 | 0.700 ± 0.074 | 3.013 ± 0.423 | 34.6 ± 3.6 | 51.5 ± 5.3 | |
| | PK-Sim | AUC | 0.575 ± 0.095 | 0.891 ± 0.099 | 3.001 ± 0.447 | 34.2 ± 1.5 | 51.0 ± 5.4 | **0.0 ± 0.0** |
| | | Cmax | 0.472 ± 0.087 | 0.930 ± 0.076 | 2.223 ± 0.168 | 46.9 ± 3.0 | 60.4 ± 3.1 | |
| Scaffold | PK-MUSE | AUC | <u>0.861 ± 0.003</u> | <u>0.534 ± 0.004</u> | **2.149 ± 0.104** | **47.0 ± 2.4** | **65.3 ± 1.4** | 1.1 ± 0.8 |
|  |  | Cmax | **0.861 ± 0.007** | **0.510 ± 0.010** | **1.818 ± 0.050** | **56.0 ± 1.0** | **76.2 ± 1.2** |  |
| | PK-MUSE<sub>Mech</sub> | AUC | 0.851 ± 0.003 | 0.552 ± 0.005 | <u>2.165 ± 0.029</u> | 45.3 ± 1.1 | <u>64.8 ± 2.4</u> | <u>1.0 ± 0.0</u> |
| | | Cmax | 0.843 ± 0.025 | 0.540 ± 0.040 | <u>1.968 ± 0.143</u> | <u>51.0 ± 4.4</u> | <u>73.5 ± 3.3</u> | |
| | HML-RF | AUC | **0.868 ± 0.012** | **0.519 ± 0.022** | 2.263 ± 0.169 | <u>46.3 ± 3.8</u> | 59.8 ± 2.6 | 8.8 ± 0.8 |
| | | Cmax | <u>0.852 ± 0.006</u> | <u>0.525 ± 0.012</u> | 1.982 ± 0.071 | 50.5 ± 2.9 | 64.2 ± 1.3 | |
| | DeepCt | AUC | 0.689 ± 0.053 | 0.795 ± 0.064 | 3.595 ± 0.425 | 28.4 ± 4.7 | 44.0 ± 3.1 | **0.0 ± 0.0** |
| | | Cmax | 0.760 ± 0.065 | 0.664 ± 0.092 | 2.951 ± 0.569 | 35.4 ± 4.9 | 53.5 ± 7.0 | |
| | PK-Sim | AUC | 0.586 ± 0.016 | 0.911 ± 0.018 | 2.693 ± 0.136 | 29.7 ± 0.8 | 54.3 ± 1.5 | **0.0 ± 0.0** |
| | | Cmax | 0.582 ± 0.015 | 0.883 ± 0.016 | 2.279 ± 0.044 | 38.2 ± 0.5 | 60.4 ± 1.2 | |
| Temporal | PK-MUSE | AUC | **0.844 ± 0.003** | **0.563 ± 0.006** | **2.012 ± 0.024** | **49.5 ± 1.4** | **67.4 ± 0.9** | <u>1.0 ± 0.0</u> |
|  |  | Cmax | **0.866 ± 0.003** | **0.510 ± 0.005** | **1.957 ± 0.117** | **52.0 ± 2.5** | **73.2 ± 1.9** |  |
| | PK-MUSE<sub>Mech</sub> | AUC | 0.824 ± 0.005 | 0.598 ± 0.009 | 2.241 ± 0.071 | 44.3 ± 1.6 | <u>66.2 ± 1.1</u> | 1.7 ± 0.8 |
| | | Cmax | <u>0.859 ± 0.008</u> | <u>0.524 ± 0.014</u> | 2.132 ± 0.200 | 48.5 ± 2.7 | <u>66.0 ± 4.1</u> | |
| | HML-RF | AUC | 0.836 ± 0.006 | <u>0.577 ± 0.011</u> | 2.252 ± 0.054 | 43.7 ± 0.7 | 61.2 ± 1.9 | 6.6 ± 0.8 |
| | | Cmax | 0.831 ± 0.008 | 0.573 ± 0.014 | <u>2.051 ± 0.125</u> | <u>48.9 ± 3.5</u> | 63.3 ± 1.3 | |
| | DeepCt | AUC | 0.693 ± 0.071 | 0.786 ± 0.089 | 3.873 ± 0.921 | 28.3 ± 2.3 | 42.7 ± 7.5 | **0.0 ± 0.0** |
| | | Cmax | 0.688 ± 0.221 | 0.744 ± 0.256 | 4.073 ± 2.790 | 35.7 ± 12.1 | 51.8 ± 17.8 | |
| | PK-Sim | AUC | 0.562 ± 0.015 | 0.943 ± 0.016 | 3.274 ± 0.538 | 33.8 ± 4.5 | 48.0 ± 3.9 | **0.0 ± 0.0** |
| | | Cmax | 0.611 ± 0.009 | 0.869 ± 0.010 | 2.507 ± 0.151 | 38.3 ± 3.5 | 58.1 ± 1.6 | |

The more telling distinction emerged as test compounds moved away from the training chemical space. On the temporal split, PK-MUSE retained an AUC MdFE of 2.012 and a *C*_max_ MdFE of 1.957, whereas DeepCt reached 3.873 and HML-RF reached 2.252 for AUC. PK-MUSE also achieved a slightly higher temporal-split AUC *R*^2^ than HML-RF (0.844 versus 0.836), together with lower RMSE, lower MdFE, and higher twofold and threefold accuracy. This stability matters because the scaffold and temporal test sets contain compounds dissimilar to those in the predefined split: mean maximum training-set Tanimoto similarity fell from 0.478 in the predefined evaluation to 0.298 in both the scaffold and temporal evaluations, and more than 90% of compounds in the latter two splits had no training compound with similarity above 0.5. Exposure performance therefore remained strong even for molecules in substantially less structurally covered regions of chemical space. PK-MUSE also maintained physiological validity under distribution shift, with invalid profiles constituting at most 1.1% of predictions across splits versus 10% for HML-RF. This gap reflects PK-MUSE’s compartmental ODE formulation, whose mass-balance constraints prevent negative concentrations and rising terminal slopes, whereas pointwise regressors offer no such trajectory-level guarantees.

### 2.3 Neural correction contributes to the out-of-distribution generalization to new chemical space and dynamic ranges

The contribution of the neural corrections became clearer when PK-MUSE was compared with its uncorrected backbone PK-MUSE_Mech_. The two models were close on the predefined split: PK-MUSE reduced AUC MdFE by 5.0% (1.794 versus 1.889), while *C*_max_ MdFE reduced by 3.0% (1.998 versus 2.060). On the scaffold split, the corrections reduced AUC MdFE by 0.7% (2.149 versus 2.165) and *C*_max_ MdFE by 7.6% (1.818 versus 1.968). The largest separation arose under temporal generalization, where PK-MUSE reduced AUC MdFE by 10.2% (2.012 versus 2.241) and *C*_max_ MdFE by 8.2% (1.957 versus 2.132). The mechanistic model is therefore already a strong structure-conditioned predictor, but the corrections add most value in the setting designed to approximate prospective prediction. Notably, the scaffold and temporal sets have almost identical mean fingerprint similarity to their training sets, yet the correction advantage is far larger on the temporal split. Chemical novelty alone thus cannot explain the difference, implying that temporal separation additionally alters the relationship between molecular structure and PK behavior. The temporal test set contains compounds with longer median infusion durations (1.0 h versus 0.5 h, range extending to 168 h) and roughly twice the median terminal half-life (6.97 h versus 2.81 h) relative to the scaffold set. These differences in disposition timescale place the temporal set in a region of PK profile space sparsely represented in training. The mechanistic model, relying on fixed compartmental parameters predicted from structure, is especially sensitive to such shifts because it cannot adapt its dynamics to unfamiliar infusion profiles or disposition timescales. The neural corrections, by contrast, adjust intercompartmental transfer and elimination rates during trajectory integration, explaining their greater benefit under temporal generalization.

The route- and phase-resolved analysis, as shown in Fig. 2, illustrates that the corrections acted non-uniformly across the concentration–time profile. We evaluated these effects using per-profile GMFE, computed from pointwise concentrations either over the full route-specific profile or within predefined trajectory phases. On the predefined split, their largest effects occurred early in the trajectory: bolus early-phase GMFE fell by 9.0% (2.002 to 1.822) and infusion input-phase GMFE by 8.2% (2.144 to 1.967), while terminal-phase performance was essentially unchanged. Under temporal generalization, this pattern shifted markedly toward late disposition: bolus terminal GMFE fell from 4.194 to 2.965 (29.3%) and infusion terminal GMFE from 3.958 to 2.812 (29.0%), with post-infusion performance also improving, though more modestly. The largest benefit of the learned dynamics therefore coincided with the region where the fixed mechanistic model deteriorated most under temporal extrapolation.

**Fig 2.**
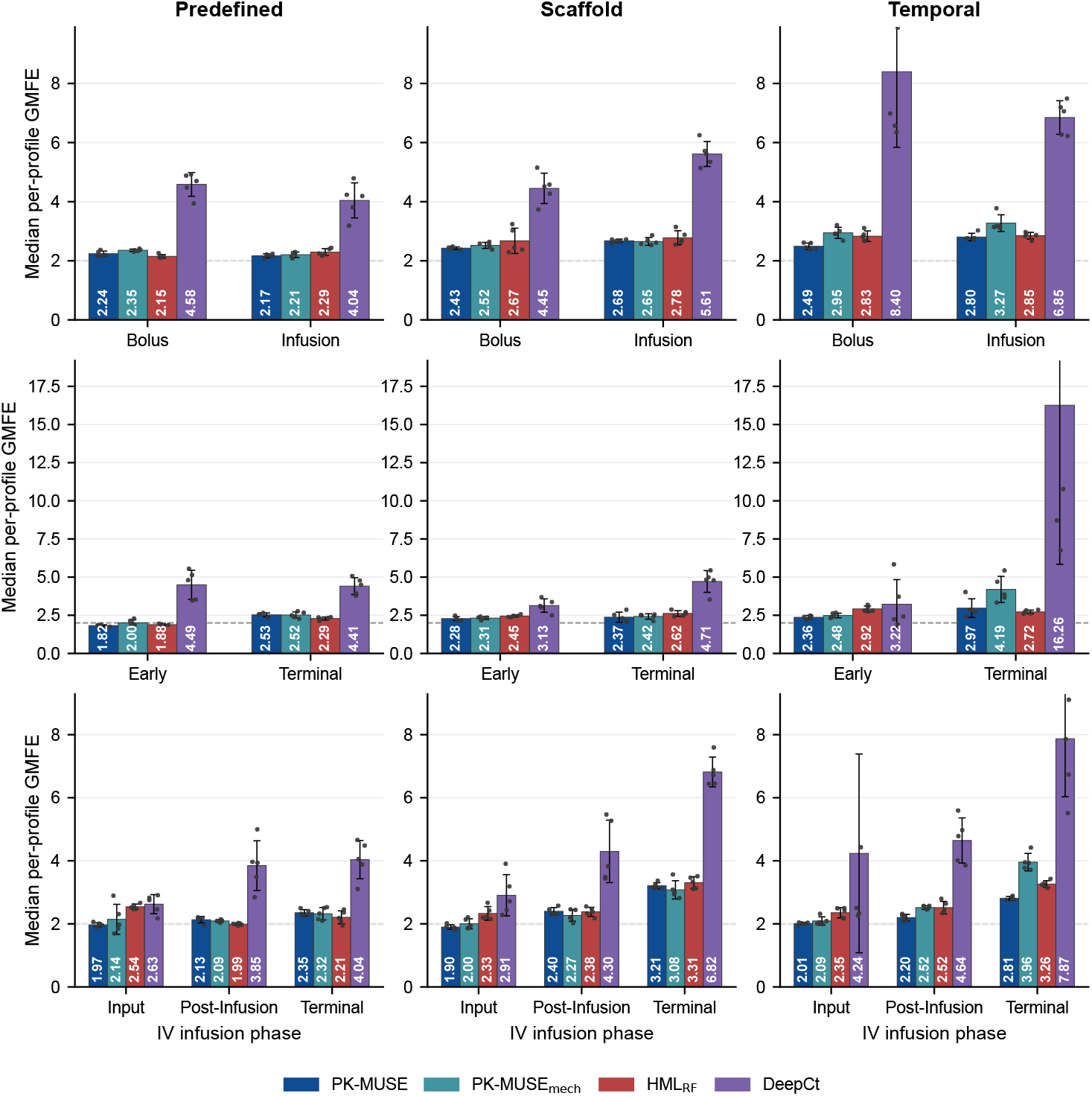
Route- and phase-specific concentration-time profile accuracy across split strategies. Bars show the median per-profile geometric mean fold error (GMFE) for PK-MUSE, PK-MUSE_Mech_, HML-RF, and DeepCt. Lower values indicate better agreement with observed concentrations. Columns correspond to the predefined, scaffold, and temporal splits. The top row stratifies complete profiles by intravenous bolus and infusion administration. The middle row partitions bolus profiles into early and terminal phases, whereas the bottom row partitions infusion profiles into input, post-infusion, and terminal phases. Per-profile GMFE was calculated from pointwise concentration fold errors within the corresponding route or phase. Error bars denote standard deviations across five runs, and points denote individual runs.

Prediction difficulty also differed by administration route. In the route-resolved analysis, PK-MUSE showed similar bolus and infusion performance on the predefined split (GMFE 2.239 versus 2.169), but infusion profiles became more difficult under scaffold and temporal generalization (2.677 versus 2.428 on scaffold; 2.804 versus 2.490 on temporal). The uncorrected backbone showed the same pattern, with infusion GMFE increasing from 2.205 on the predefined split to 2.654 on scaffold and 3.274 on temporal. Within infusion profiles, error concentrated after drug input rather than uniformly: PK-MUSE GMFE rose from 1.967 during infusion to 2.132 post-infusion and 2.354 during terminal disposition on the predefined split, and from 2.011 to 2.200 and 2.812 under temporal generalization. The learned corrections produced their largest infusion-phase benefit in late disposition, reducing temporal-split terminal-phase infusion GMFE by 29.0% (3.958 to 2.812), compared with only 3.9% during active infusion (2.093 to 2.011). Together, these results show that PK-MUSE improves concentration-time prediction most strongly in late-disposition regions, especially under temporal generalization.

### 2.4 Learned corrections localize mechanistic misspecification

PK-MUSE’s learned corrections were highly asymmetric across the three kinetic processes (Fig. 3A). Averaged across splits and seeds, the mean absolute corrections to central-to-peripheral and peripheral-to-central transfer were |*ϵ*_12_| = 0.254 and |*ϵ*_21_| = 0.223, compared with only |*ϵ*_el_| = 0.100 for elimination. The distribution corrections were also more prevalent: *ϵ*_12_ and *ϵ*_21_ exceeded an absolute magnitude of 0.1 at 86% and 79% of time points, versus 44.5% for elimination. At the compound level, 31% of molecules required almost no elimination correction (|*ϵ*_el_| *<* 0.05), compared with only 2-3% for the distribution corrections. The corrections were further phase-dependent, being largest and most sign-consistent during terminal disposition, where on the temporal split *ϵ*_12_ averaged +0.35 and *ϵ*_21_ averaged −0.30, versus +0.16 for *ϵ*_el_. This pattern indicates that the learned dynamics accelerate central-to-peripheral transfer while slowing peripheral-to-central return late in the profile, effectively deepening the peripheral compartment during terminal disposition, precisely where the fixed two-compartment model deteriorates most. The temporal split consistently required the largest corrections across all three terms, consistent with its greater mechanistic misspecification.

**Fig 3.**
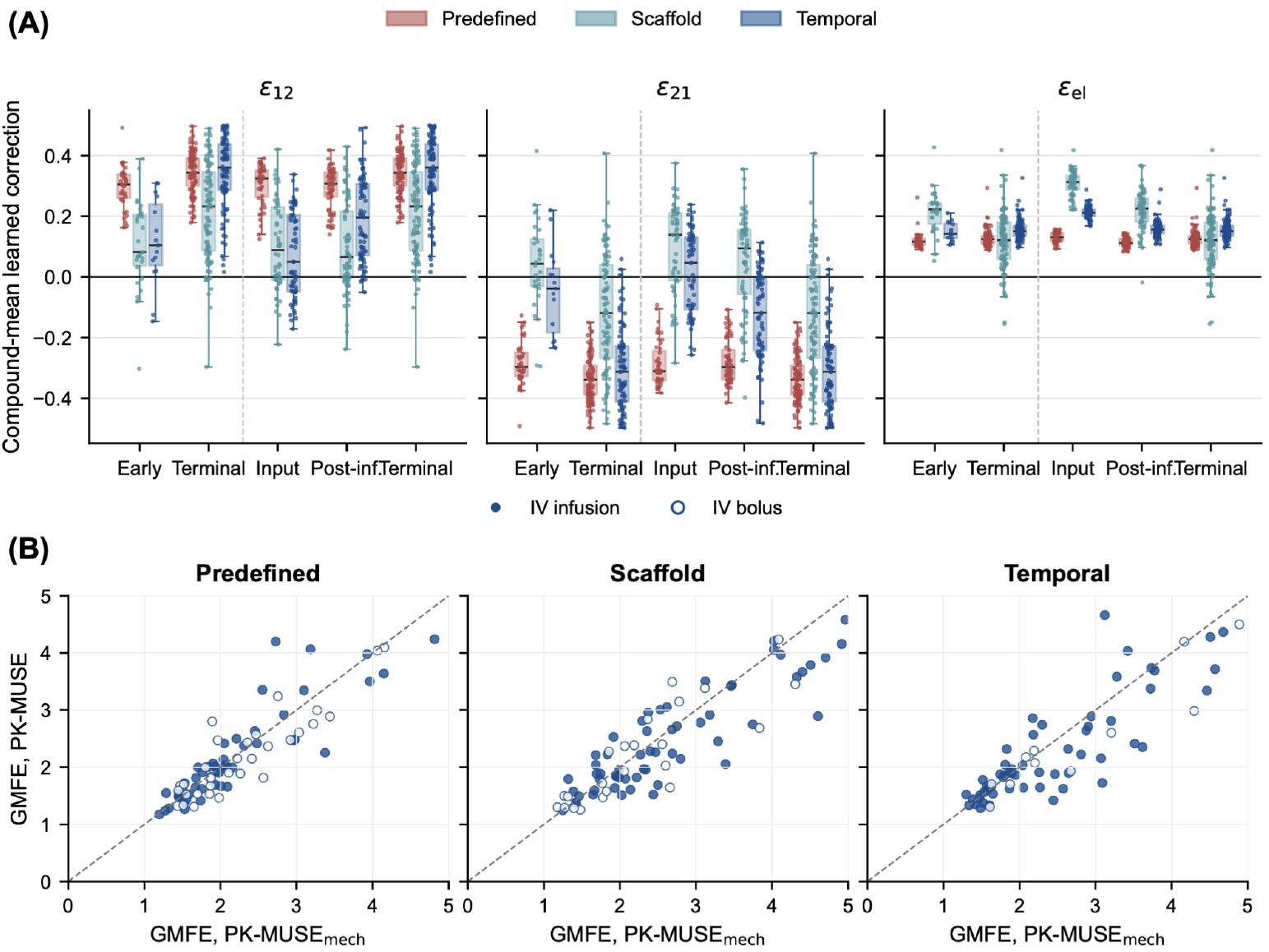
Learned corrections and their effect on per-compound prediction error. **A**, Compound-mean learned corrections to the three kinetic processes (*ϵ*_12_, central-to-peripheral transfer, *ϵ*_21_, peripheral-to-central transfer, and *ϵ*_el_, elimination) by profile phase and split.**B**, Per-compound GMFE of PK-MUSE versus its mechanistic backbone (PK-MUSE_Mech_) for each split. Points below the identity line indicate compounds improved by the learned corrections. The corrections preferentially rescue the compounds with the largest mechanistic error, most strongly on the temporal split.

To establish where the corrections help, we compared per-compound prediction error (via per-compound GMFE) between PK-MUSE and its mechanistic backbone directly (Fig. 3B). The corrections improved accuracy for 59% of compounds in each split, but their benefit concentrated on the compounds the mechanistic model predicted worst. On the temporal split, compounds with mechanistic GMFE above 3 improved in 64% of cases (Δ log_10_ GMFE = −0.043), versus 54% (−0.003) for those already within threefold.

The corrections thus act primarily as a rescue mechanism for mechanistic failures under distribution shift rather than as a uniform refinement of adequate predictions. A minority of compounds moved above the identity line, indicating that the corrections degraded an otherwise adequate mechanistic prediction; these correction-associated failures are analyzed below.

The property dependence of this benefit further localized the misspecification (Fig. 4). On the temporal split, the corrections helped most for compounds at the extremes of the training property distribution: low clearance (Δ log_10_ GMFE ≈ −0.7), long terminal half-life (≈ −1.0), low fraction unbound (≈ −0.55 for highly protein-bound compounds), and extreme acidic or basic *pK*_*a*_. Trends on the predefined and scaffold splits were largely flat, indicating that this property-dependent benefit is specific to temporal generalization rather than a generic feature of the correction network. The pattern is consistent with the error-correlation analysis, in which clearance showed the strongest relationship with per-compound GMFE (Pearson *r* = − 0.55 on log-clearance for PK-MUSE, predefined split), terminal half-life correlated moderately with mechanistic error under temporal generalization (*r* = 0.38), and fraction unbound correlated negatively with temporal-split error (Spearman *ρ* = −0.42). Together, these results indicate that the fixed two-compartment model is most misspecified for compounds at sparsely represented extremes of the property space, and that the learned corrections partially compensate, which is why their benefit concentrates where the mechanistic model is weakest.

**Fig 4.**
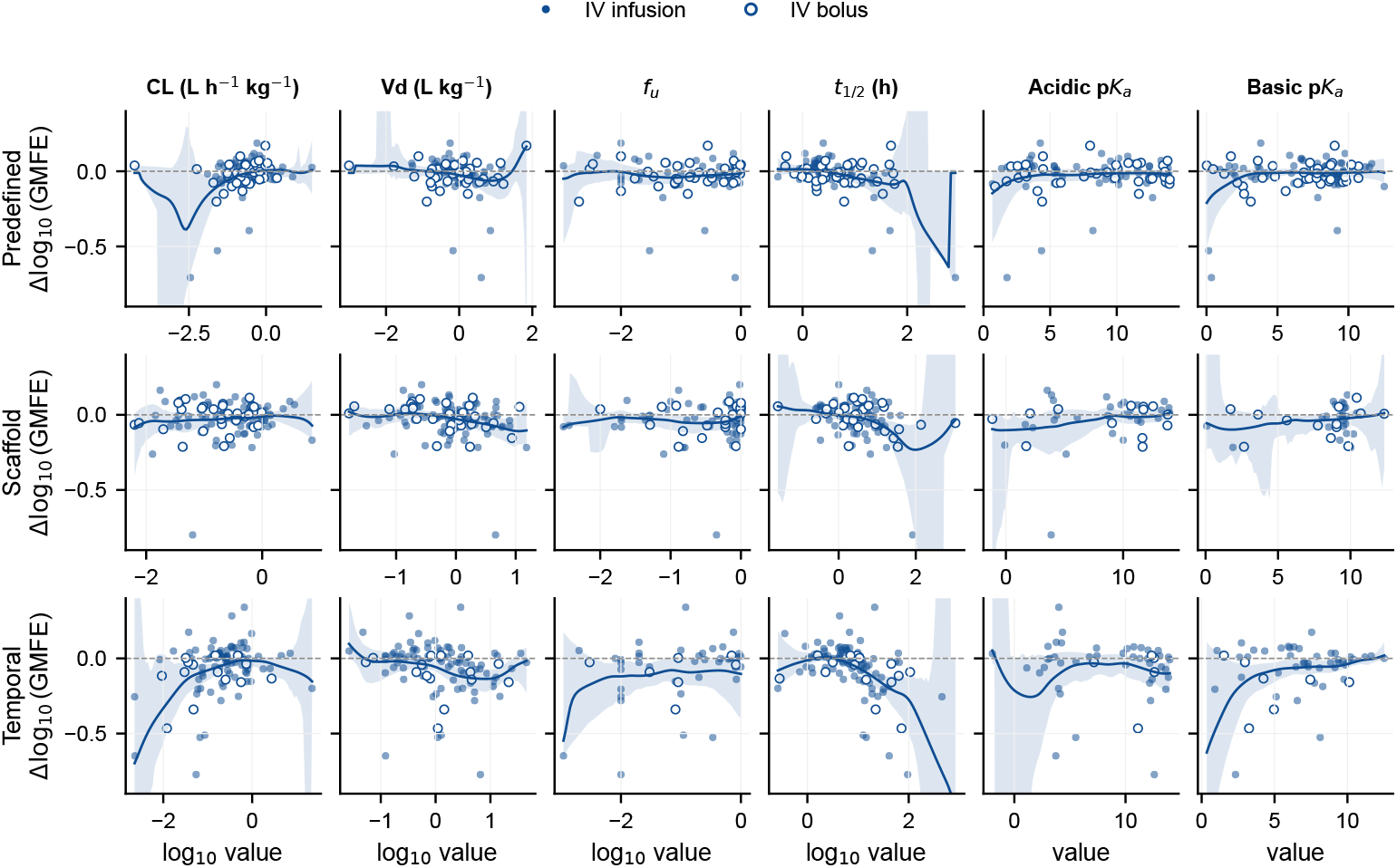
Property dependence of the correction benefit. Each panel shows the change in per-compound log_10_ GMFE between PK-MUSE and PK-MUSE_Mech_ 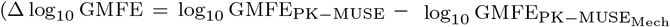. Negative values indicate improvement by the corrections) as a function of six compound pharmacokinetic properties (clearance, volume of distribution, fraction unbound, terminal half-life, and acidic and basic *pK*_*a*_), for each split. Filled and open points denote intravenous infusion and bolus profiles, respectively. The corrections confer the largest benefit at property extremes, low clearance, long terminal half-life, low fraction unbound, and extreme *pK*_*a*_.

### 2.5 Prediction failures concentrate in early-phase misspecification

Although PK-MUSE improved aggregate accuracy, its errors concentrated in a relatively small subset of compounds. We defined a prediction failure as a per-compound concentration GMFE greater than 3, together with physiologically invalid profiles. PK-MUSE failed for 32.3%, 41.5%, and 44.9% of compounds in the predefined, scaffold, and temporal splits. It nonetheless maintained physiological validity throughout, with invalid profiles constituting at most 0.6% of predictions across splits (Table 1), so the failure burden is almost entirely accuracy-limited rather than validity-limited.

To locate where these failures arise, we partitioned each observed profile into early (*t* ≤ *t*_1*/*2_) and terminal (*t > t*_1*/*2_) phases using the observed terminal half-life and computed phase-specific GMFE, flagging a compound as early-phase-misspecified when its early-phase GMFE was ≥ 2. Among PK-MUSE’s valid failures at GMFE *>* 3, this flag was carried by 96.4%, 84.0%, and 89.4% of failing compounds across the predefined, scaffold, and temporal splits. The continuous early-phase GMFE classified overall failure (GMFE *>* 3) with an AUC of 0.88 pooled across splits and seeds, confirming that the dominant failure mode lies in the early phase rather than in terminal disposition.

A second contributor to failure was extreme PK parameters, such as very short or long terminal half-lives, high or low clearance, or unusually large or small volumes of distribution, accounting for 56.4%, 75.8%, and 59.7% of valid failures across splits. However, extreme PK parameters rarely caused failure independently of early-phase misspecification: failures that were extreme-PK but not early-phase were only 3.0%, 15.5%, and 7.5%, whereas those that were both extreme-PK and early-phase reached 53.4%, 60.3%, and 52.2%. A substantial fraction were early-phase without extreme PK (43.0%, 23.7%, and 37.1%), and almost none fell in neither category (0.6%, 0.5%, and 3.1%). Extreme PK parameters therefore act as a mediated risk factor, raising the likelihood of early-phase misspecification, the proximate cause of failure, rather than degrading prediction independently. Dopamine illustrates this: with a terminal half-life of 0.34 h and clearance of 7.24 L/hr/kg, both dataset extremes, its GMFE improved from 10.37 with PK-MUSE_Mech_ to 6.44 with PK-MUSE, yet the residual early-phase error kept it in the failure set.

This early-phase concentration of failure was not specific to PK-MUSE. Among valid failures at GMFE *>* 3, the early-phase flag was carried by 84-96% of failing compounds across all four models and all three splits, pointing to a shared limitation of structure-to-PK frameworks rather than a deficiency unique to the compartmental ODE approach. The models diverged instead on the accuracy-validity trade-off. PK-MUSE and PK-MUSE_Mech_ kept *>*99% of profiles physiologically valid (0.2-0.8% invalid), whereas HML-RF produced 6.8-9.0% invalid profiles with negative concentrations, rising terminal slopes, or sustained post-infusion rises that violate mass balance. DeepCt produced no invalid profiles but failed for 66.5-77.7% of compounds at GMFE *>* 3, more than double PK-MUSE’s rate. PK-MUSE thus occupied the most favorable position: near-zero invalidity with the lowest failure rate among mechanistically valid models.

## 3 Discussion

PK-MUSE demonstrates that a UDE framework combining a two-compartment pharmacokinetic backbone with bounded neural corrections can predict human intravenous concentration–time profiles from chemical structure while retaining mechanistic interpretability. Across random, scaffold, and temporal splits, PK-MUSE consistently outperformed pointwise machine learning, deep learning compartmental, and PBPK baselines. Because the corrections are interpretable adjustments to named pharmacokinetic processes, elimination, central-to-peripheral distribution, and peripheral-to-central redistribution, their magnitudes reveal that misspecification in the fixed two-compartment model concentrates in intercompartmental distribution, particularly during terminal disposition.

The dominance of distribution corrections over the elimination correction has implications for how compartmental models are specified. Classical two-compartment models treat distribution as a first-order process with fixed rate constants, but the learned corrections suggest that effective distribution rates vary along the trajectory in ways a fixed-parameter model cannot capture. This aligns with the literature on compartmental misspecification, where inadequate representation of distribution phases biases terminal-phase estimates and derived parameters such as clearance [11]. By letting state-dependent neural corrections adapt these rates, PK-MUSE captures such behavior without adding compartments and the accompanying model complexity.

The temporal split provides the most informative test of the framework. Although the scaffold and temporal test sets share nearly identical mean Tanimoto similarity to their training sets, the correction advantage is far larger under temporal generalization (10.2% AUC MdFE reduction versus 0.7% for scaffold). We attribute this to differences in route composition and disposition timescale: the temporal set contains more infusion compounds with longer infusion durations and terminal half-lives. The mechanistic model, predicting fixed compartmental parameters from structure, cannot adapt to these shifts, whereas the neural corrections adjust intercompartmental transfer rates during trajectory integration, explaining their disproportionate terminal-phase benefit under temporal generalization.

Several limitations should be noted. First, PK-MUSE was evaluated only on intravenous data. Oral absorption introduces additional complexity, including absorption kinetics, first-pass metabolism, and transit times, that the current framework does not address. Second, the dataset comprises 718 compounds, large for PK profile prediction but small relative to standard molecular machine learning benchmarks, and, like other machine learning models, the framework should improve with more data. Third, the neural corrections can occasionally degrade an otherwise adequate mechanistic prediction, particularly for compounds at the extremes of the pharmacokinetic parameter space. Finally, PK-MUSE is currently limited to two-compartment systems; however, this framework can be extended to multi-compartment pharmacokinetic models.

The USE (Universal System Equations) formulation used here is general and extensible. Adding an absorption compartment with a learned input function would enable oral PK prediction. Additionally, incorporating patient-level covariates into the molecular representation could support population PK modeling. Coupling the PK model to a pharmacodynamic model through the learned concentration trajectory could enable integrated PK/PD prediction. More broadly, retaining a mechanistic backbone while learning bounded, interpretable corrections to specific rate processes offers a template for combining domain knowledge with data-driven flexibility across pharmacological modeling.

## 4 Methods

This section describes the problem formulation, PK-MUSE framework in detail, the experimental setup consisting of the dataset and baselines, and the evaluation metrics.

### 4.1 Problem formulation

For a compound with Simplified Molecular Input Line Entry System (SMILES) string *S* administered intravenously at dose *D*, we predict its plasma concentration–time trajectory *C*(*t*) under either bolus or finite-duration infusion dosing. The model receives the molecular SMILES, dose, infusion duration and time grid, and predicts plasma concentrations at the corresponding time points.

### 4.2 PK-MUSE Architecture

#### 4.2.1 Chemical descriptor-derived pharmacokinetic representation

PK-MUSE (Stage-2) uses a pharmacokinetically defined molecular representation. In the first stage, molecular structure is mapped to compound-level pharmacokinetic and physicochemical quantities using molecular representations derived from Molecular ACCess System (MACCS) keys, Functional-Class Fingerprints 6 (FCFP6) fingerprints, RDKit descriptors and Mordred descriptors to predict clearance (CL), volume of distribution (*V*_*D*_), plasma fraction unbound (*f*_*u*_), mean residence time (MRT), acidic *pK*_*a*_ and basic *pK*_*a*_. The resulting six-dimensional vector

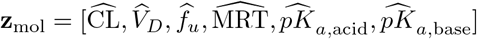

provides the molecular input to the dynamical model. This was inspired by the hierarchical machine-learning framework of Jia et al. [6].

#### 4.2.2 Molecule-conditioned mechanistic backbone

The mechanistic component of PK-MUSE is a classical two-compartment intravenous pharmacokinetic model, following the central–peripheral compartmental formulation established in early pharmacokinetic theory [25, 26]. In this model, drug in the bloodstream is represented by a central compartment. The drug can move from the central compartment into a peripheral compartment, which represents tissues where distribution occurs more slowly, and can later return to the central compartment. Drug elimination occurs from the central compartment. The two compartments are mathematical representations of kinetically distinct spaces rather than anatomical organs.

We formulate the system in terms of drug amounts. Let *A*_1_(*t*) and *A*_2_(*t*) denote the amounts of drug in the central and peripheral compartments, respectively. Their corresponding concentrations are:

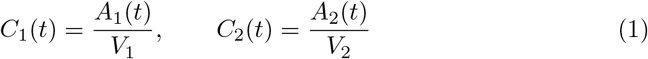

where *V*_1_ and *V*_2_ are the apparent central and peripheral volumes of distribution. For an intravenous input *R*(*t*), the mechanistic dynamics are:

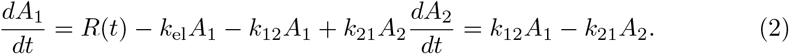

Here, *k*_el_ is the first-order elimination rate from the central compartment, *k*_12_ refers to the distribution from the central to the peripheral compartment, and *k*_21_ refers to the redistribution from the peripheral to the central compartment. The observed plasma concentration is (*C*_1_(*t*)). This formulation separates three processes that determine the shape of an intravenous concentration–time profile: elimination from the systemic circulation, outward tissue distribution, and subsequent redistribution back to the central compartment.

PK-MUSE makes this conventional mechanistic system compound specific by conditioning its parameters on the molecular pharmacokinetic representation **z**_mol_. Four multilayer perceptrons map **z**_mol_ to the base elimination rate *k*_el_, central volume *V*_1_, intercompartmental clearance *Q*, and ratio of *V*_2_*/V*_1_. The intercompartmental rate constants are then derived as:

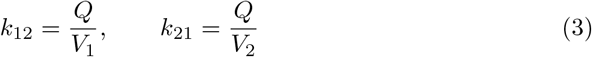

Thus, *Q* defines the common exchange capacity between the two compartments, while the compartment volumes determine the corresponding first-order transfer rates. Positive output transformations are applied to *k*_el_, *V*_1_, *Q*, and *V*_2_, ensuring positive base rates and volumes. We additionally learn *k*_el_ directly from **z**_mol_.

The initial conditions of the compartmental system are derived using the molecular pharmacokinetic representation. The peripheral compartment starts drug-free, *A*_2_(0) = 0. For the central compartment, PK-MUSE uses the classical bolus initial condition *C*_1_(0) = *D/V*_ic_, where *V*_ic_ is an initial-concentration volume set to either the network-learned central volume *V*_1_ described above, or the Stage-1 predicted volume of distribution 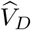. We saw better performance when using *V*_ic_ = *V*_1_, since that is the same learned volume which scales the compartment’s concentration and anchors its initial condition. Deriving *V*_ic_ instead from the Stage-1 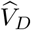 introduced invalid trajectories (1.0%) and the worst *C*_max_ MdFE among the ablations.

The administered dose should enter the trajectory only once, either through the initial condition or through the infusion input *u*(*t*). PK-MUSE therefore sets the initial condition according to the administration route:

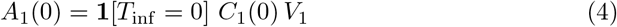

where *T*_inf_ denotes the infusion duration. For a bolus dose (*T*_inf_ = 0), the dose is introduced through the nonzero initial condition. For infusions with finite duration (*T*_inf_ *>* 0), the initial condition is set to zero and the dose is delivered through *u*(*t*) over the infusion period. This distinction is necessary to preserve mass balance: without it, an infusion dose would be counted twice, once in the initial condition and again through *u*(*t*). Consistent with this, removing the gate worsened performance for both exposure endpoints, yielding MdFEs of 1.95 for AUC and 2.12 for *C*_max_, compared with the best-performing PK-MUSE model.

#### 4.2.3 State-dependent neural rate corrections

PK-MUSE extends PK-MUSE_Mech_ by allowing the elimination and intercompartmental distribution rates to change as the pharmacokinetic trajectory evolves. PK-MUSE retains the mechanistic equations and learns bounded corrections to the three rate constants *k*_el_, *k*_12_, and *k*_21_ as opposed to learning a vector field without the mechanistic structure support. A shared neural correction network receives the compound-level pharmacokinetic representation **z**_mol_, a scaled representation of time, the current central and peripheral states, and the infusion status. The input vector is:

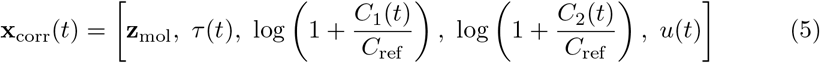

Where

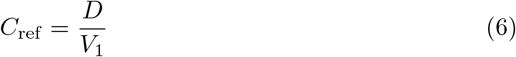

is the bolus-equivalent concentration associated with the administered dose. Scaling *C*_1_ and *C*_2_ by *C*_ref_ expresses the current state relative to the compound’s dose-dependent concentration scale, while the log(1 + *x*) transformation compresses the wide range of concentrations encountered over a PK trajectory. Time is represented relative to the compound-specific elimination timescale as:

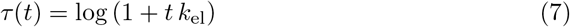

Thus, the correction network receives elapsed time relative to the expected disposition rate of the compound rather than absolute time alone. The correction network consists of a shared multilayer perceptron with two hidden layers and three outputs,

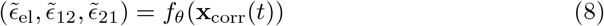

Each output is transformed to a bounded log-rate correction,

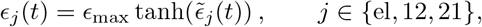

with *ϵ*_max_ = 0.5. The effective rates used during ODE integration are

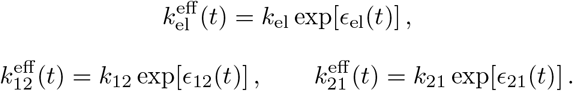

The bound |*ϵ*_*j*_ |≤ 0.5 limits each instantaneous rate modification to between *e*^*−*0.5^ ≈ 0.61 and *e*^0.5^ ≈ 1.65 times its corresponding mechanistic rate. The output layer of the correction network is initialized to zero, such that all three corrections are initially zero and PK-MUSE begins training with dynamics identical to PK-MUSE_Mech_.

### 4.3 Experimental setup

#### 4.3.1 Dataset

Our training consists of two stages similar to the hierarchical apporach by Jia et al. [6], both the datasets for prediction of PK parameters as well as prediction of PK-profiles for model training, validation, and independent testing was extracted from the human intravenous PK profile study from the same. We selected this dataset because it links human concentration-time profiles with compound-level PK annotations and therefore supports benchmarking and analysis of profile prediction models. For concentration-time profile prediction, plasma concentration-time profiles were manually extracted from published figures with WebPlotDigitizer [27]. The primary sources were those referenced in the curated human PK database of Lombardo et al. [28]. This design maintains consistency between the extracted profiles and the corresponding parameter estimates. Eligible profiles came from healthy adult subjects who received a single intravenous dose. The curation retained both intravenous bolus and intravenous infusion profiles. For compounds investigated at multiple clinical dose levels, profiles were preferentially selected from dose regimens approximating 1 mg/kg [6]. Data curation and quality-assurance procedures followed the high-throughput PBK/PBPK methodologies described by Geci et al. and Komissarov et al. [29, 30]. Stage-1 data consists of For the stage-2 training we ensured that there was no overlap of training data from stage-1. In this study, each data instance is one compound-level intravenous concentration-time profile. The model input includes the compound SMILES string, dose, infusion duration, initial concentration estimate, and observed sampling times. The model output is the predicted plasma concentration at each observed time point. Concentrations are represented in ng/mL, time in hours, dose in mg/kg, and infusion duration in minutes before conversion to hours. The final dataset contains profiles for 717 unique compounds and 8,707 concentration observations. We excluded profiles with two or fewer concentration observations before model training. We aggregated duplicate measurements at the same compound time point with the median concentration. We removed extreme concentration values to reduce digitization or reporting artifacts. We log-transformed concentrations for model training because concentration values span several orders of magnitude. The models were tested on three splits - the predefined split, scaffold split, and temporal split. We refer to the dataset split used by Jia et al. [6] as the predefined split as used in their studies, the compounds in the test set of the predefined split were compounds with known experimental PK parameter values through literature, hence the name predefined [6, 28]. It contains 613 training compounds and 104 independent test compounds, this is the same split that has been used in our work. In the work, the predefined and random splits are used inter-changably. Additionally, we evaluated the models on a scaffold and temporal-based splits. The scaffold split tests generalization to unseen chemical scaffolds. The temporal split separates compounds by year-of-disclosure metadata, which ranged from 1878 to 2014. A cutoff of 1997 separated earlier compounds from later compounds. All the splits had experiments repeated over five random seeds. The final reported values for the splits are the mean and standard deviation across five runs. For the predefined, and time split the seed impacted the initialization and validation set creation because the train and test compounds were fixed, whereas for the scaffold split the different scaffold groups were shuffled for each seed.

#### 4.3.2 Baseline models

We compared PK-MUSE with three baselines: PK-Sim [31], HML-RF [6], and DeepCt [32].

PK-Sim served as the non-ML mechanistic baseline. We ran whole-body PBPK simulations in PK-Sim version 12.2. We used one standardized baseline human template for all compounds, so simulation differences mainly reflected compound-specific inputs. The compound-specific inputs to PK-Sim differed by split strategy. For the predefined split, all inputs were experimentally measured values available from the Lombardo et al. database [28]: molecular weight, log *P*, *f*_*u*_, acidic and basic *pK*_*a*_ values, CL, infusion duration, and halogen count. For the scaffold and temporal splits, experimental PK parameters were not uniformly available for the test compounds, because those compounds were not selected on the basis of parameter availability. Fraction unbound (*f*_*u*_) and lipophilicity (log *P*) were therefore predicted using the trained random forest models from chemical descriptors, and the simulation model’s input was correspondingly reduced by omitting the halogen count and *pK*_*a*_. We represented administration as IV bolus or IV infusion according to the reported infusion duration. For tissue distribution, we did not commit to a single partition-coefficient model. Instead, we ran each simulation independently under five tissue-distribution methods implemented in PK-Sim/ospsuite: PK-Sim Standard, Berezhkovskiy, Poulin and Theil, Rodgers and Rowland, and Schmitt. We report the mean and standard deviation of each exposure metric across these five methods, analogous to the mean and standard deviation reported across five random seeds for the machine-learning baselines, so that PK-Sim’s reported spread reflects sensitivity to the choice of tissue-distribution model rather than random variation. We automated simulations with the ospsuite package in R version 4.5.2.

HML-RF follows the hierarchical machine-learning framework introduced by Jia et al. [6]. In this framework, QSAR models first predict compound-level physicochemical and PK parameters from chemical structure using multiple fingerprint-based representations of the chemical structure. A second-stage random forest model then predicts concentration values at individual time points. The features include *f*_*u*_, CL, *V*_Dss_, acidic and basic *pK*_*a*_ values, time, and infusion duration. Unlike PK-MUSE, HML-RF does not model the concentration trajectory as the solution of an ODE. It predicts each concentration value directly from the feature vector.

DeepCt served as the deep-learning mechanistic-compartmental baseline. DeepCt predicts positive conventional compartmental parameters from molecular features, including clearance, central volume, intercompartmental clearance, and peripheral volume terms [32]. These parameters are converted into a compartmental rate matrix, and concentration-time profiles are generated with a fixed analytical linear-system solution for bolus or infusion dosing. The original DeepCt implementation used MEL-LODDY embeddings from a cross-pharma federated learning model [33]. Because those embeddings are proprietary, we retrained DeepCt with RDkit fingerprints.

Two recent deep-learning pharmacokinetic models could not be evaluated on this dataset. Uni-PK [34] embeds a neural ODE within a mechanistically grounded structure and was developed and evaluated primarily on rat concentration–time data. When we attempted training Uni-PK on this human intravenous dataset diverged through numerical overflow: with explicit integration and a priori unbounded learned rates, unstable trial steps produce states that grow without bound within a single solve, after which non-finite losses and gradients corrupt the model parameters and the predicted initial conditions of subsequent iterations. We were unable to obtain converged Uni-PK models and therefore do not report metrics for it. LSTM-PK [18] predicts concentration sequences with a long short-term memory network trained on simulated profiles sampled on uniform time grids. As its authors note, ‘the LSTM architecture used in our model requires uniformly spaced time steps so extending the model to handle irregularly sampled PK data would require further methodological adaptation’ [18]. The present dataset consists of irregularly sampled clinical profiles whose sampling grids differ across compounds, so LSTM-PK cannot be applied without such adaptation.

#### 4.3.3 Evaluation metrics

We evaluated predictive performance at two complementary levels: concentration-time profile accuracy and exposure-endpoint accuracy. We additionally assessed generalization beyond the training compounds and physiological validity. Each test instance comprised one intravenous concentration-time profile for one compound.

Profile accuracy was evaluated from the agreement between predicted and observed concentrations at the observed sampling timepoints. Exposure-endpoint accuracy was evaluated using area under the concentration-time curve (AUC), which summarizes systemic exposure, and *C*_max_, which captures peak exposure. For each compound, observed and predicted AUC and *C*_max_ values were derived independently from the corresponding concentration-time profiles. AUC was calculated from the first to the last observed sampling timepoint using the same numerical integration procedure for observed and predicted profiles. *C*_max_ was defined as the maximum concentration at the observed sampling timepoints.

Because plasma concentrations and pharmacokinetic endpoints span several orders of magnitude, raw-scale absolute errors can overemphasize compounds with large concentration or exposure values. We therefore evaluated prediction errors in logarithmic and fold-error space [6, 18, 32].

##### Concentration-time profile accuracy

For a generic observed quantity *y* and corresponding prediction directional fold error as *ŷ*, we defined

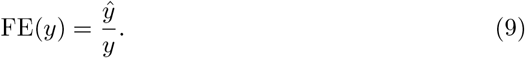

For compound *j* and observed sampling timepoint *i*, concentration fold error was therefore FE(*C*_*ij*_) = *Ĉ*_*ij*_ */C*_*ij*_, where *C*_*ij*_ and *Ĉ*_*ij*_ denote the observed and predicted concentrations, respectively. Pointwise concentration errors were aggregated within each compound to obtain a profile-level geometric mean fold error,

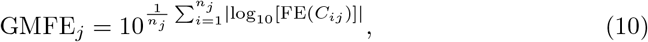

where *n*_*j*_ is the number of observed sampling timepoints for compound *j*. GMFE has an optimum of 1 and treats equivalent magnitudes of underprediction and overprediction symmetrically.

Profile-level error was also summarized by root mean squared error in log-concentration space,

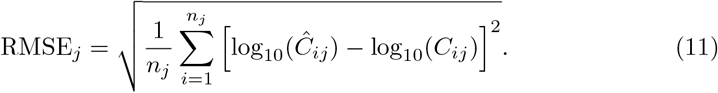

GMFE and RMSE quantify agreement across the complete observed concentration-time trajectory rather than agreement in a derived exposure endpoint. Unless otherwise specified, GMFE refers to the per-profile concentration-time GMFE: point-wise concentration fold errors are aggregated across the observed sampling timepoints within each compound profile. This quantity measures concentration-time accuracy and is distinct from the MdFE reported for the derived AUC and *C*_max_ endpoints.

##### Exposure-endpoint accuracy

For an exposure endpoint *Y*_*j*_, where *Y*_*j*_ denotes either AUC or *C*_max_, endpoint fold error was calculated using the same definition, FE(*Y*_*j*_) = *ŷ*_*j*_ */Y*_*j*_, where *Y*_*j*_ and *ŷ*_*j*_ are the observed and predicted endpoint values, respectively. The magnitude of endpoint error was evaluated in absolute logarithmic space,

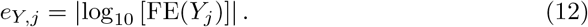

Across compounds, median fold error was calculated as

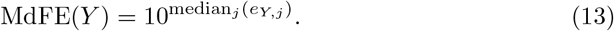

MdFE(*Y*) therefore represents the typical multiplicative error in prediction of AUC or *C*_max_.

Twofold accuracy was defined as the percentage of compounds satisfying

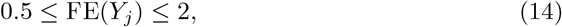

and threefold accuracy was defined analogously as the percentage satisfying

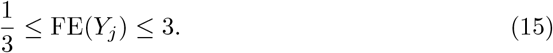

Agreement between observed and predicted exposure endpoints across compounds was additionally characterized using the coefficient of determination, *R*^2^, calculated on log-transformed endpoint values:

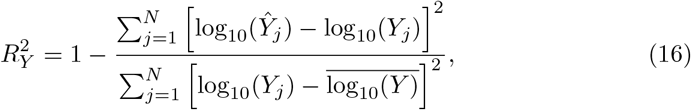

where *N* is the number of compounds and 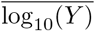 is the mean log-transformed observed endpoint value across compounds.

Together, 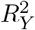, MdFE(*Y*), and twofold and threefold accuracy characterize complementary aspects of endpoint prediction: agreement in variation across compounds, typical multiplicative error, and the proportions of compounds predicted within predefined fold-error ranges.

##### Handling of physiologically invalid profiles

Numerical agreement with observed concentrations or exposure endpoints does not necessarily imply that a predicted concentration-time trajectory is physiologically valid. A predicted profile was classified as invalid if it exhibited one or more of the following behaviors: (i) one or more negative predicted plasma concentrations; (ii) a positive predicted log-concentration slope during the observed terminal phase; or (iii), for infusion profiles, a sustained increase in predicted concentration after the infusion had ended. Negative plasma concentrations are physiologically impossible [35], while terminal concentrations are expected to decline during elimination [10, 35].

Invalid profiles were retained in the evaluation rather than removed. For fold-error-based metrics, an invalid profile, or a profile for which a finite positive concentration or endpoint prediction could not be calculated, was assigned

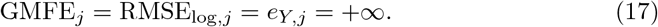

Consequently, invalid compounds remained in the denominator for median fold-error summaries and were treated as outside the twofold and threefold accuracy ranges. The frequency of invalid profiles was additionally reported as a separate metric. End-point *R*^2^ was calculated across compounds with finite positive observed and predicted endpoint values.

##### Route- and phase-specific profile accuracy

Aggregate AUC and *C*_max_ errors summarize overall exposure agreement but do not identify where along the concentration-time trajectory prediction errors occur. This distinction is particularly relevant for intravenous data because bolus and infusion dosing have different input dynamics. Following an IV bolus, the early profile reflects rapid distribution after essentially instantaneous input, whereas an infusion profile contains a period of continuous drug input followed by post-infusion distribution and terminal elimination. Classical PK models therefore treat bolus and infusion as distinct input conditions [10].

We first stratified test compounds by administration mode and evaluated IV bolus and IV infusion profiles separately. This analysis assessed whether aggregate performance was driven preferentially by one route and whether a model represented instantaneous and finite-duration drug input equally well.

We subsequently partitioned each observed profile into pharmacokinetic temporal phases. Phase boundaries were determined from the observed profile and then applied identically to predictions from all models. For IV bolus profiles, observations at or before the compound-specific median observed sampling time were assigned to the early phase, and later observations were assigned to the terminal phase. For IV infusion profiles, observations collected during infusion were assigned to the input phase. Post-infusion observations were then divided at the compound-specific median post-infusion sampling time into post-infusion and terminal phases.

Within each route or phase, profile accuracy was calculated using the same concentration-level GMFE definition described above, with the timepoint index restricted to the corresponding subset. This decomposition complements exposure-endpoint metrics because similar overall AUC values can arise despite compensating prediction errors at different portions of the concentration-time trajectory. Phase-specific profile errors therefore assess whether a model accurately represents drug input, distribution, and terminal disposition rather than only their aggregate effect on exposure.

## Funding

This work was supported by R01GM122845 (NIGMS of NIH), R01AG057555 (NIA of NIH), R33AG083302 (NIA of NIH), and R41DA062978 (NIDA of NIH).

## Author Contributions

A. B. conceived the concept, designed the method and the experiments, prepared data, implemented the algorithms, performed the experiments, analyzed results, and wrote the manuscript. L. C. prepared data and performed the experiments. M. K. performed the experiments. J. W. conceived the concept and acquired funding. L. X. conceived the concept, designed the method and the experiments, wrote the manuscript, and acquired funding.

## Competing interests

The authors declare no competing interests.

## Notes

### Competing Interest Statement

The authors have declared no competing interest.

